# Long-term soil fungal community recovery after fire is impacted by climate change

**DOI:** 10.1101/2020.10.09.333443

**Authors:** Spencer McGee, Alyssa Tidwell, Erin Riggs, Hannah Veltkamp, Geoffrey Zahn

## Abstract

Though much is known about fungal importance to forest health, there is very little information about factors that impact soil fungal community recovery time after a fire. Soil samples were taken from burn sites within one ecotype of temperate coniferous forest in Utah over a 20-year chronosequence. Sites were selected from available historic burns and were similar in plant community structure, elevation, slope, and aspect. Fungal DNA from these samples was compared to soil from paired unburned sites nearby to measure community similarity and estimate soil fungal recovery rates. Differences between paired burned and unburned sites remained fairly stable over a decadal time scale overall, but fungal community structure was found to recover more quickly in areas with a higher average annual temperatures. A significant positive correlation in community recovery was seen in areas with a difference of as little as two degrees celsius per year. The only other environmental variable that significantly interacted with time since burn was annual mean precipitation. As global temperatures increase, alpine fires are increasing as well, but these results suggest that fungal community recovery time will be shortened under new climate scenarios.

## Introduction

The effect of fires on belowground microbial communities has received increased attention in the past decade due to the importance of microbes to overall ecosystem recovery. Soil microbes interact with plants during disturbance recovery to shape resource availability, community assembly and successional trajectories (Knelman et al., 2015; Sikes et al., 2016; Weidner et al., 2015). Ecosystem recovery from fire disturbance is thus dependent on microbes that facilitate plant succession either directly via obligate symbioses (Dickie et al., 2017) or indirectly via processes such as resource liberation and soil stabilization (Bonner et al., 2019; Claridge et al., 2009; Fuentes-Ramirez et al., 2018).

Currently, the reported effects of fire on belowground microbial communities have few consistencies other than a general agreement that full recovery to pre-fire conditions takes place on a decadal time scale (Holden et al., 2013; Köster et al., 2014; Treseder et al., 2004). Reported differences between studies include varied recovery times for bacteria and fungi (Bárcenas-Moreno et al., 2011; Bárcenas-Moreno & Bååth, 2009) and divergent effects of fire on microbial diversity and abundance in differing biomes (Allison et al., 2010; Dove & Hart, 2017; Hansen et al., 2019). The majority of studies have focused on time since fire as the main effect in their particular ecosystems, and the contrasting details about microbial recovery could be specific to the climatic and edaphic properties of the biomes in which they were observed.

Warmer and drier conditions in many ecosystems due to climate change are increasing the frequency and impact, and decreasing the predictability of, wild fires (Abatzoglou & Williams, 2016; Flannigan et al., 2009). Thus, it is important to continue building knowledge of how soil microbes respond to fires. Furthermore, it is important to understand not only how future climate scenarios alter fire regimes, but whether they will alter the recovery responses of microbial communities in those systems.

Here, we conducted an observational experiment along a 20-year fire chronosequence, comparing paired burned and unburned sites along an annual mean temperature gradient in an alpine ecosystem dominated by pinyon pine and juniper forests in Utah, USA. This ecosystem is expected to continue warming at a faster rate than most other biomes with a 20-year projected climate that includes an annual mean temperature of 2 deg C above baseline according to the IPCC AR5 regional synthesis report (Barros et al., 2014). The study design allowed us to test the hypothesis that a warmer climate could alter belowground microbial recovery rates after fires within a single biome.

## Methods

### Overview

A range of historic fire events was used to select 11 forested regions of Utah that had burned between 1 and 20 years prior to our study. Each burned site was paired with an adjacent unburned site to serve as controls, giving us a total of 22 sites. Sites were selected to maximize similarity across a range of environmental variables. At each site, we collected burned and unburned soil from the organic horizon of 3 locations, measured canopy and ground cover vegetation, and collected vegetation samples for identification and herbarium curation. We used meta-amplicon sequencing to characterize fungal community structure from each soil sample.

### Site Selection

The Utah Department of Natural Resources sourced Utah fire data for the previous 20 years. 300 GPS points 100 m inside of burn boundaries were used to compile national GIS layers including topography, land use and cover, human impacts, and climate data. Habitats were dominated by Pinyon Pine and Juniper, with common understory plant communities dominated by Asteraceae and Poaceae. Sites were spread across Utah from 38.5 to 41.6 deg latitude and ranged in elevation between 1483m and 2969m, with a median elevation of 1660m (Fig. 1). The annual mean temperature at sites ranged from −7.9C to −3.4C with a median of −5.0C. Soil types were predominantly a mixture Nielsen, Sterling, and Steed series.

**Fig. 1.**
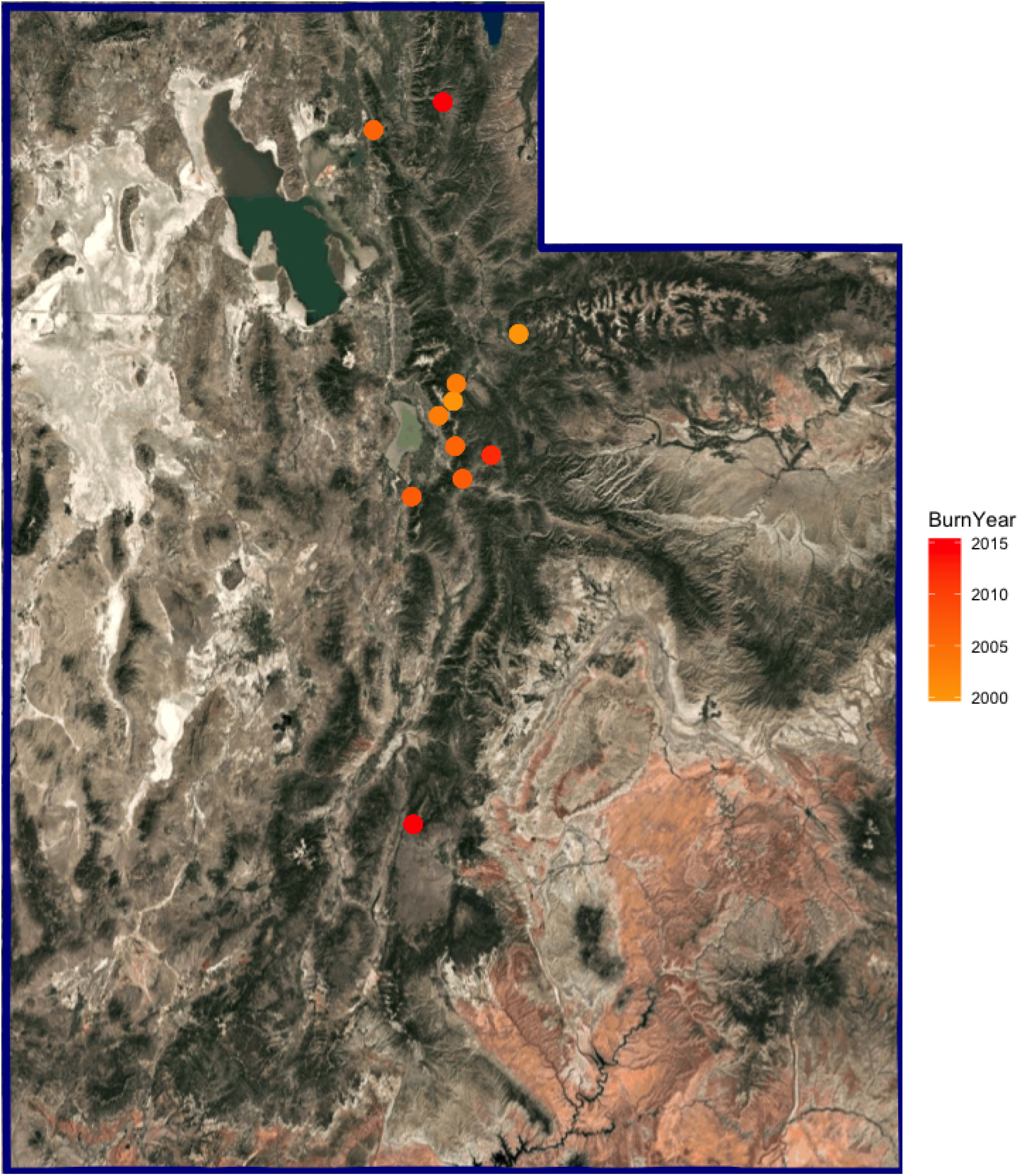
Map of site locations, colored by burn year, Each point represents an adjacent paired burned and unburned site.

Using these and other site characteristics (see S.I. Table 1), we built a pairwise distance matrix for every possible combination of variables, and selected a subset of the 300 sites that had minimal land variation, but still maintained substantial ranges in burn chronology and annual mean temperature (see S.I. Fig. 1). All raw site data and full code for selecting study sites can be found in the Supporting Information.

### Field Collection

Photography, site marking, and sample collection protocols were designed before field work began. Three soil samples were collected with sterile utensils from each site at a depth of 5 to 10 cm 0.5m from the site center in the North, East, and West cardinal directions. Soils were frozen at −80° within 6 hours of collection. Local plant cover was recorded and photographed and representative plant samples were collected for later identification. International Field Collection Forms were filed for each plant collection and all specimens are housed in the Utah Valley University Herbarium (UVSC).

### DNA Extraction and Library Preparation

Genomic DNA from 0.25g of soil was extracted from each sample using DNeasy PowerSoil Kits (QIAGEN, Venlo, The Netherlands). Fungal DNA was amplified with the ITS1F (CTTGGTCATTTAGAGGAAGTAA; Gardes & Bruns 1993) and ITS2 (GCTGCGTTCTTCATCGATGC; White et al., 1990) primers, modified with the addition of Illumina adaptors (Caporaso et al., 2011) using the following protocol: 98 2 min; 22 cycles of: 98 15 s, 52 30 s, 72 30 s; 72 2 min). After 22 cycles, the PCR product was diluted 1:12 and 1 μL of this was used as a template for 8 more rounds of PCR with a 60 deg annealing temperature in which bi-directional barcodes bound to reverse complemented Illumina adaptors acted as primers. Resulting barcoded libraries were cleaned, normalized, and sequenced with the Illumina MiSeq platform (V3 chemistry, 2 × 300 bp).

### Bioinformatics

Reads were demultiplexed and barcode sequences were removed by the sequencing center. Quality filtration and bioinformatics were performed in R. Briefly, we extracted the ITS1 region, filtered forward reads based on quality, utilized a clustering-free Divisive Amplicon Denoising Algorithm (DADA) to infer Amplicon Sequence Variants (ASVs) (Callahan et al., 2016), removed chimeras and potential contaminants and assigned taxonomy against a custom ITS1 database.

The ITS1 region of the rDNA was extracted from all raw reads using ITSxpress (Rivers et al., 2018). Quality control on ITSxpress output consisted of removing reads with ambiguous base calls and those with a maxEE of >2, and truncating each read when quality scores dropped
 below 20. Due to lower quality, and to reduce false-positive detection, reverse reads were not used (Pauvert et al., 2019). Filtered forward reads were subjected to *de novo* chimera detection and removal in DADA2 and potential contaminants were inferred from extraction negatives and removed from all samples using the prevalence method in the *decontam* package (Davis et al., 2018). Cleaned and filtered ASVs were assigned taxonomy with the RDP Classifier algorithm against a custom database consisting of the UNITE database (v. 1.12.2017) and a custom set of outgroups including ITS1 sequences from metazoans and viridiplantae taken from NCBI. The outgroups added to UNITE can be found in the Supporting Material. Any sequences matching non-fungal taxa were removed. The remaining ASVs that were unambiguously assigned to fungi were used in all downstream analyses within the*phyloseq* R package (McMurdie & Holmes, 2013). All sequences have been deposited in the NCBI Sequence Read Archive under the accession PRJNA550446.

### Statistical Analyses

All analyses were performed in R (Version 3.4.4). A PermANOVA model of community composition as an interactive function of Burn Year, Fire Treatment, and Location was fit using the adonis function of the *vegan* package (Okansen et al., 2016). A community distance matrix was generated with the vegdist function of the *vegan* package. The community distance between paired burned and unburned sites at the same location was regressed against time since burn and decadal annual mean temperature in a linear mixed-effect model using the *lme4* package (Bates et al., 2015) with paired community distance as a response and annual mean temperature and years since burn as predictors. P-values were obtained using the *lmerTest* package (Kuznetsova et al., 2017). Differential abundance of taxa between burn treatments was analyzed using a beta-binomial model with the *corncob* R package (Martin et al., 2020).

## Results

No fungal community of any burned site we observed had fully rebounded to the alpha diversity or community structure of its unburned counterpart within the 20 year timeframe that was studied. All samples were dominated by Ascomycota and Basidiomycota, regardless of whether they were burned or not, but the relative dominance of these two main phyla decreased in older sites. In burned sites 14 years or older, Ascomycota and Basidiomycota still dominated in terms of relative abundance, but other phyla such as Mucormycota and Olpidiomycota were also more common (Fig. 2). Relative abundance of taxonomic groups did not correlate significantly with annual mean temperature, elevation, latitude, or soil type.

**Fig. 2.**
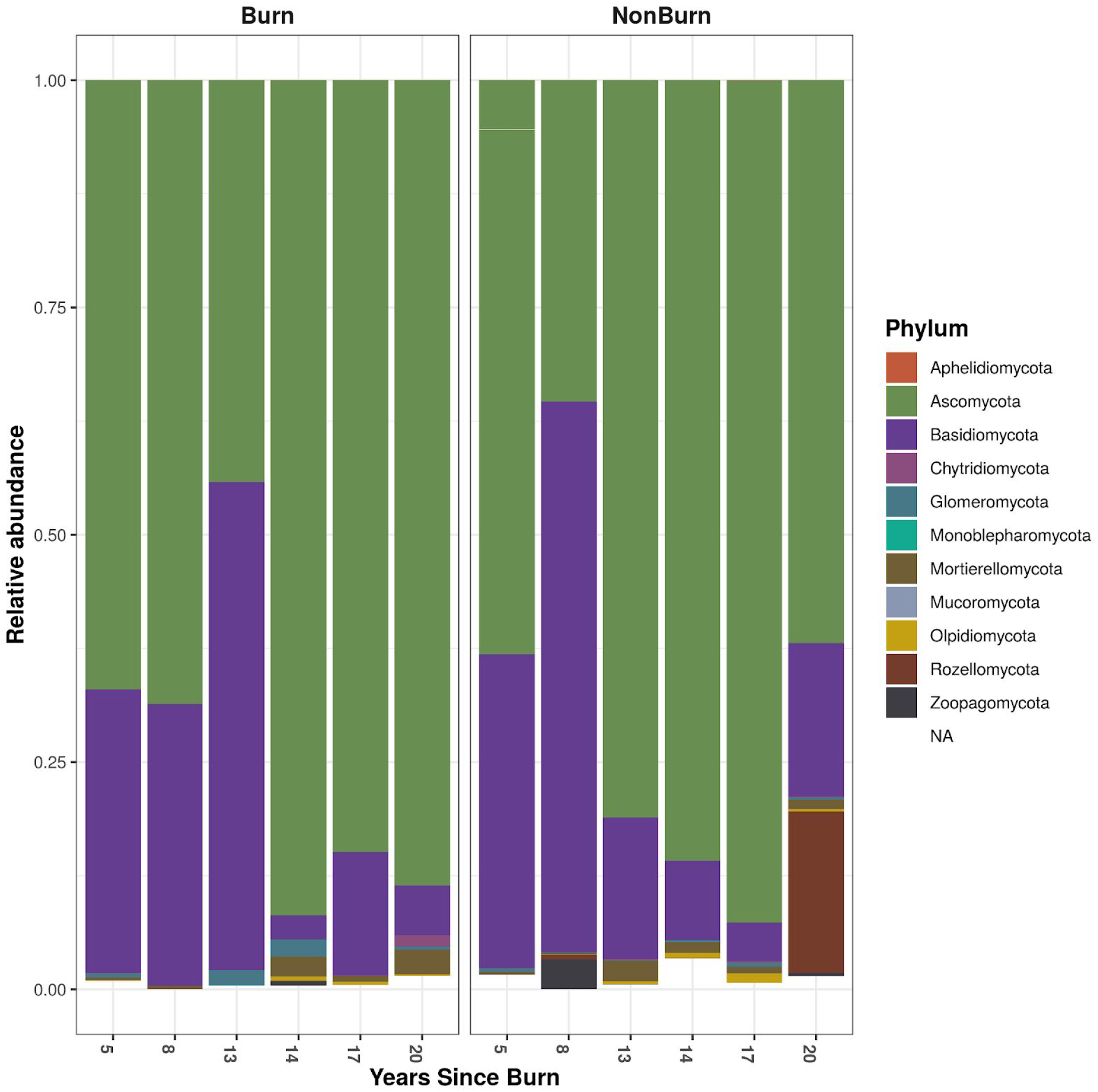
Stacked barplot of phylum relative abundance. Samples were grouped by burn chronology.

Alpha diversity was not significantly different overall between aggregated burned and unburned sites (Fig. 3), and there was no consistent pattern between burn status and Shannon diversity in between-site pairs.

**Fig. 3.**
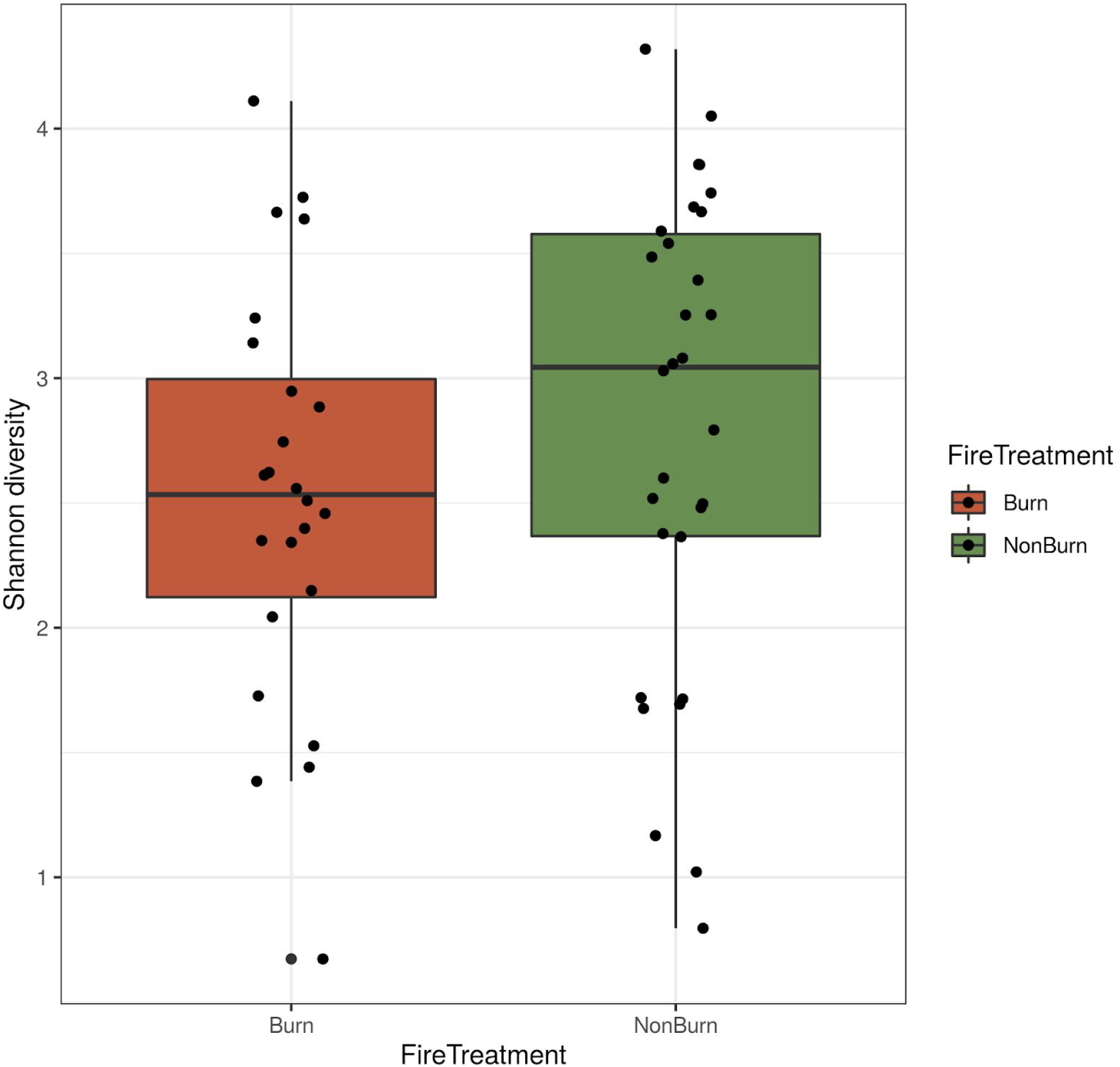
Shannon diversity by fire treatment

Beta diversity was significantly affected by burn status (P = 0.035), time since burn (P=0.001), and annual mean temperature (P=0.001) in an interactive PermANOVA model (S.I. Table 2). Further, there was a significant interaction term between time since burn and annual mean temperature (P=0.002).

Mean community distance between paired burned and unburned sites showed a small but significant decrease over time since burn, congruent with previous research showing the decadal scale of recovery for fungi after fires. However, in this study, we also found that annual mean temperature was an important predictor of the rate of recovery (Fig. 4; LMER; P < 0.0005), and each temperature increase of 1 deg C accounted for an additional 5% community similarity along our full chrono-series (S.I. Table 3).

**Fig 4.**
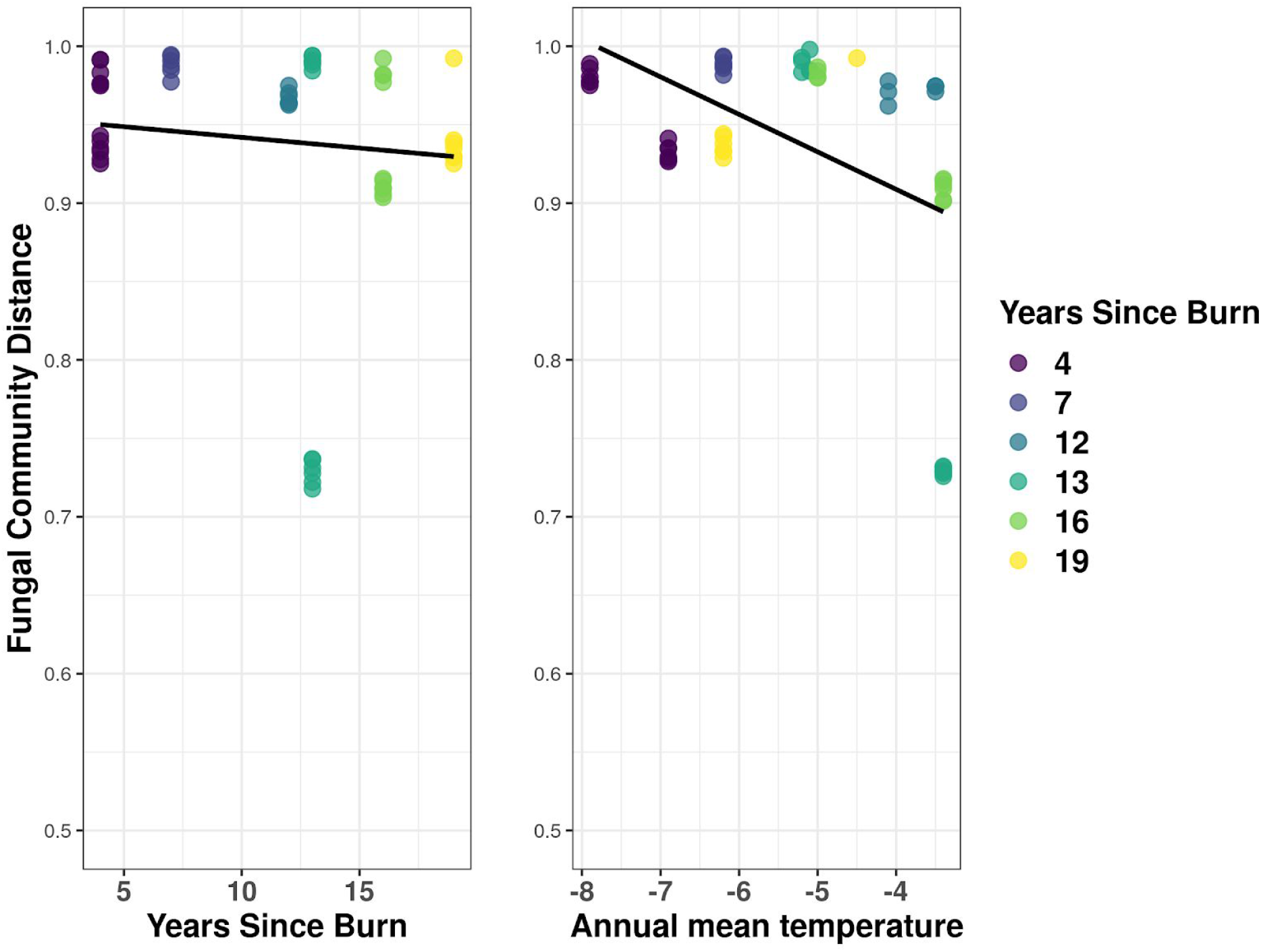
Effect of annual mean temperature on community distance between paired adjacent burned/unburned sites. Community distance of 1 Indicates no overlap between fungal community structure between paired burned and unburned sites. Warmer temperatures Ced to quicker recovery

When location-specific plant community data was added to linear mixed-effect models, no significant correlation between plant cover or plant community and paired fungal community distance was found.

Differential abundance analyses yielded 4 fungal families that were significantly enriched in either burned or unburned locations across all sites (Fig. 5). The basidiomycetes Serendipitaceae and Thelophoraceae had greater abundance and variability in unburned sites, and the ascomycetes Microascaceae and an unclassified family-level taxon in Dothidiales had greater abundance and variability in burned sites.

**(SI?)Fig. 5.**
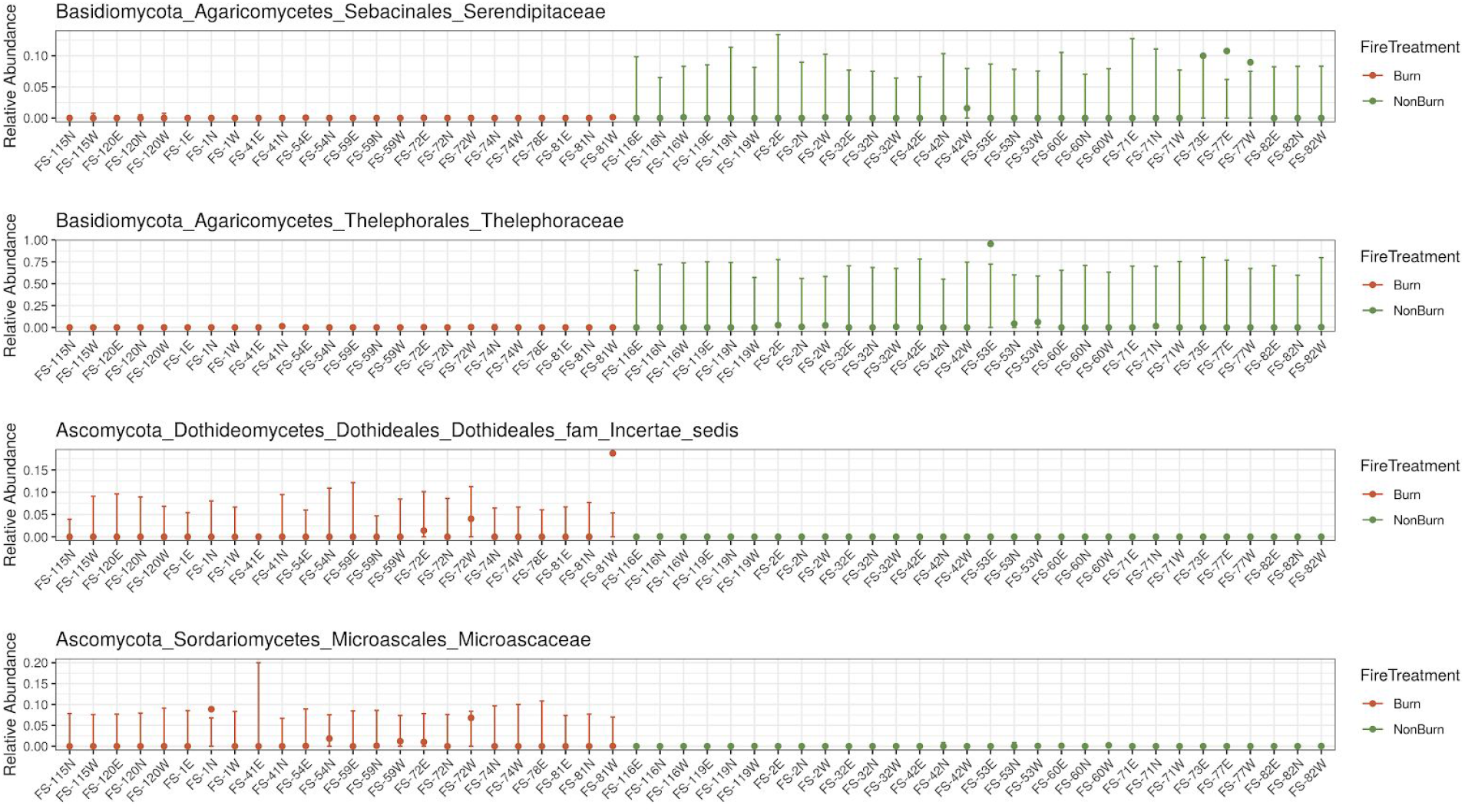
Fungal taxa at the family-level with significant differential abundance between burn treatments. Beta-binomial model; Wald test P < 0.05

## Discussion

Though most studies on fungal recovery after fires have taken place in a single biome and focused on chronology as the main predictor, Dove and Hart (2017) showed in a meta-analysis that the magnitude of at least the initial fungal community responses to fire can vary based on fire frequency and biome type. These are important observations because they suggest hypotheses to help explain observed discrepancies in fungal recovery trajectories between studies.

Here, we tested whether large-scale climatic factors could also help explain variance within a single biome. Comparing nearby pairs of burned and unburned sites along a forest fire chronosequence showed that climatic conditions can have a role in soil fungal community recovery trajectories. In particular, our observation that annual mean temperature of a site has a small but significant effect on recovery speed is a possible example of how local environmental variation can help explain discrepancies in fungal recovery times after fires.

Due to ongoing recent climate change, any chronosequence studies must inevitably deal with the potential covariance of time since burn and mean annual temperature. There was some correlation between these two variables in the present study, with older sites tending toward lower mean annual temperatures, which could be an effect of climate change. We were unable, however, to acquire site-specific environmental variables from 20 years prior, to test this due to lower resolution of these observations from that time period.

Additionally, climate warming is inextricably linked to other variables that have been shown to affect fire severity and fire frequency such as reduced soil moisture and increased fuel load (Harvey, 2016). These factors could also be important drivers of fungal recovery trajectories. These legacy factors were not considered in the present study since none of our selected sites had burned in the previous 75 years and were from very similar biomes, but they may be important to consider in work of this nature.

Our results indicate that even within a single biome, slight alterations to climatic variables like mean annual temperature can affect the rate of fungal recovery from fires. This has implications for broader disturbance modeling efforts. For example, the reduction and rebounding of soil fungal communities is important for dynamic vegetation models, but many disturbance models use predictors that are simulated in isolation, neglecting any potential interaction effects (Seidl et al., 2017). As research into fungal recovery from fires moves forward, we need to take note of site-level climatic conditions in addition to other variables of interest.

## Supporting Material

### Supporting figures

**S.I. Fig 1.**
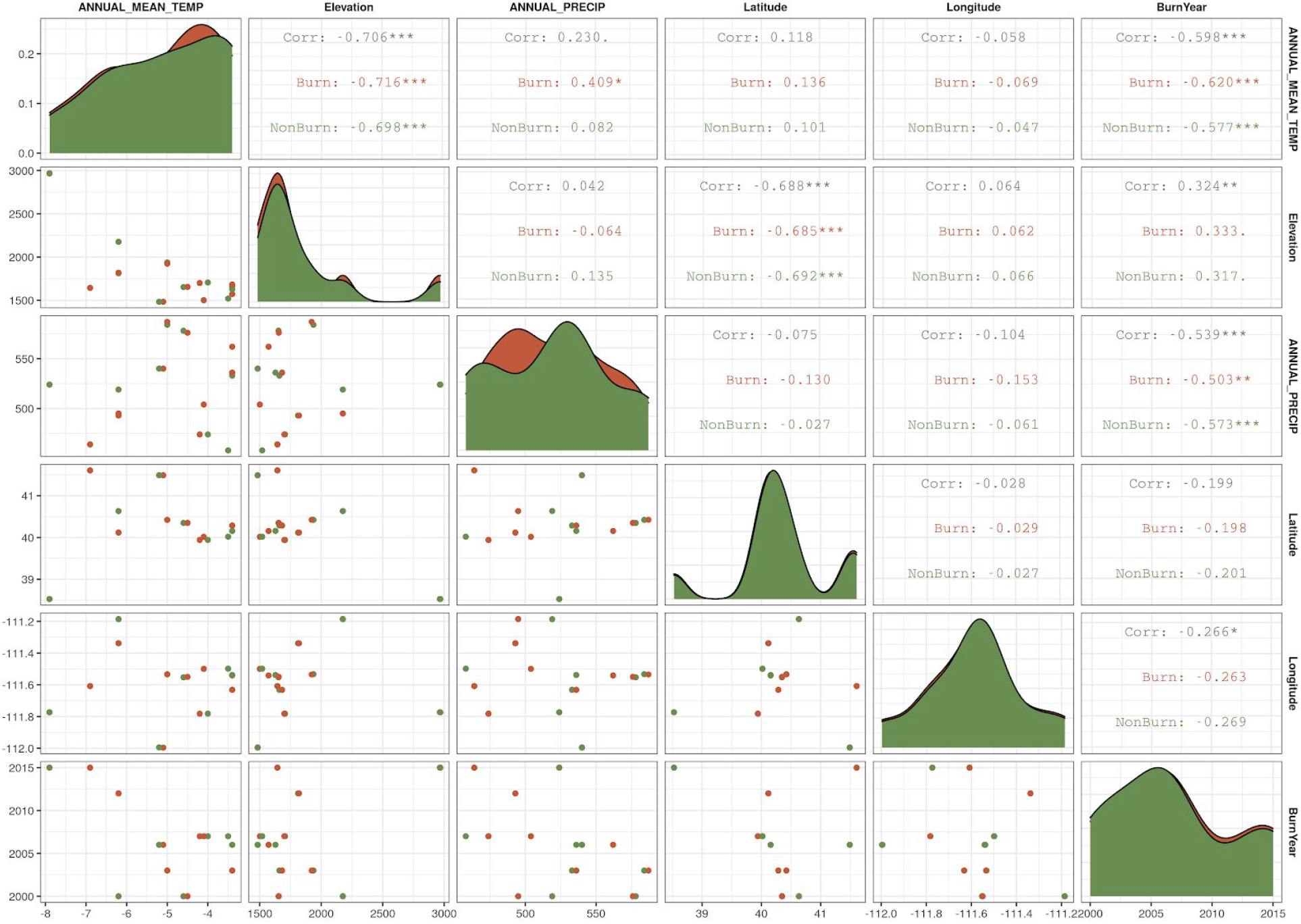
Correlation plot between paired-site variables

**S.I. Fig. 2.**
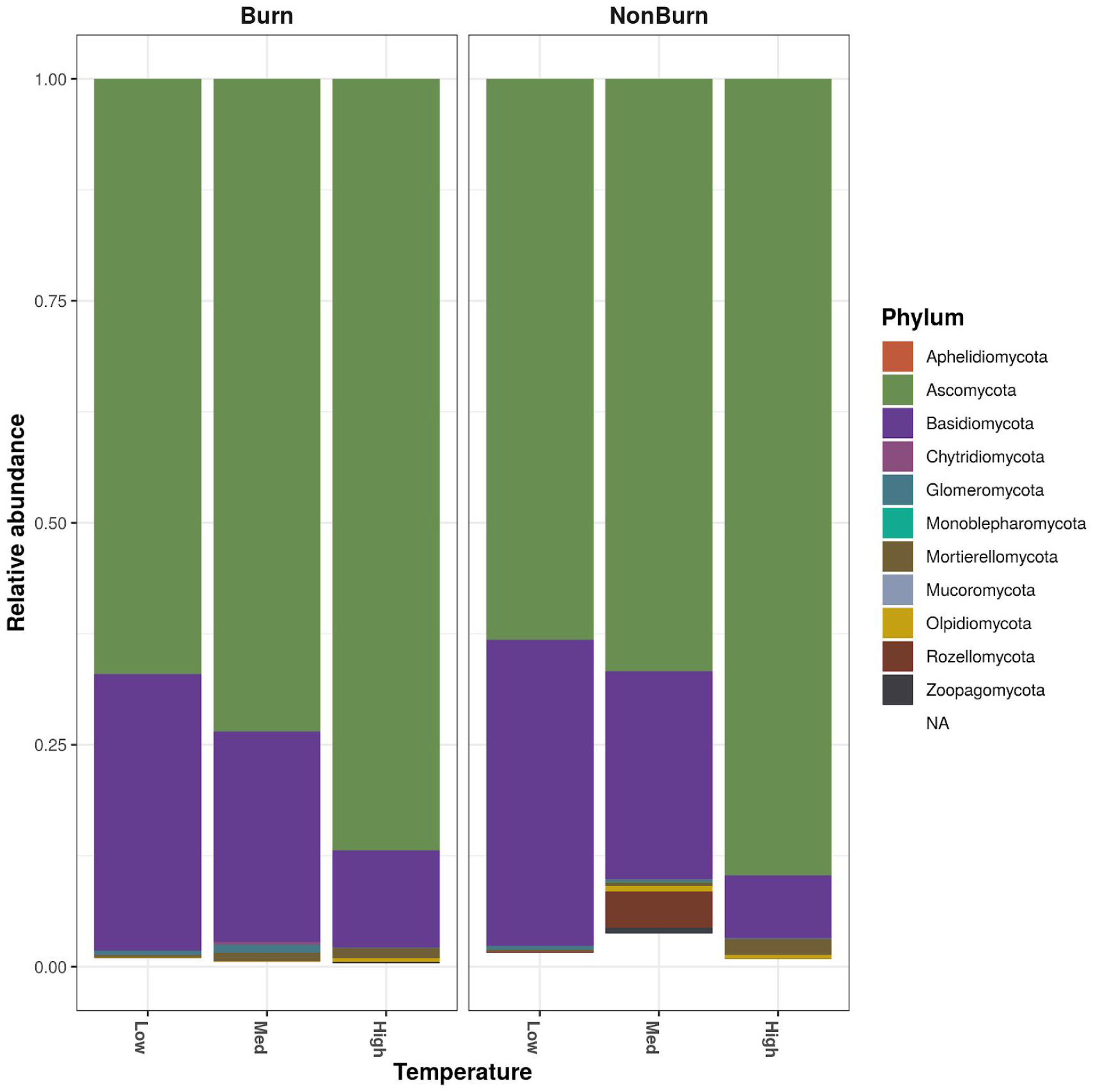
Fungal phylum relative abundance based on Annual Mean Temperature groups. Grouped as follows: “Low” = −7.9 > x < −6.4; “Med” = −6.4 > x < −4.9; “High” = −4.9 > x < −3.4

**S.I. Fig. 3.**
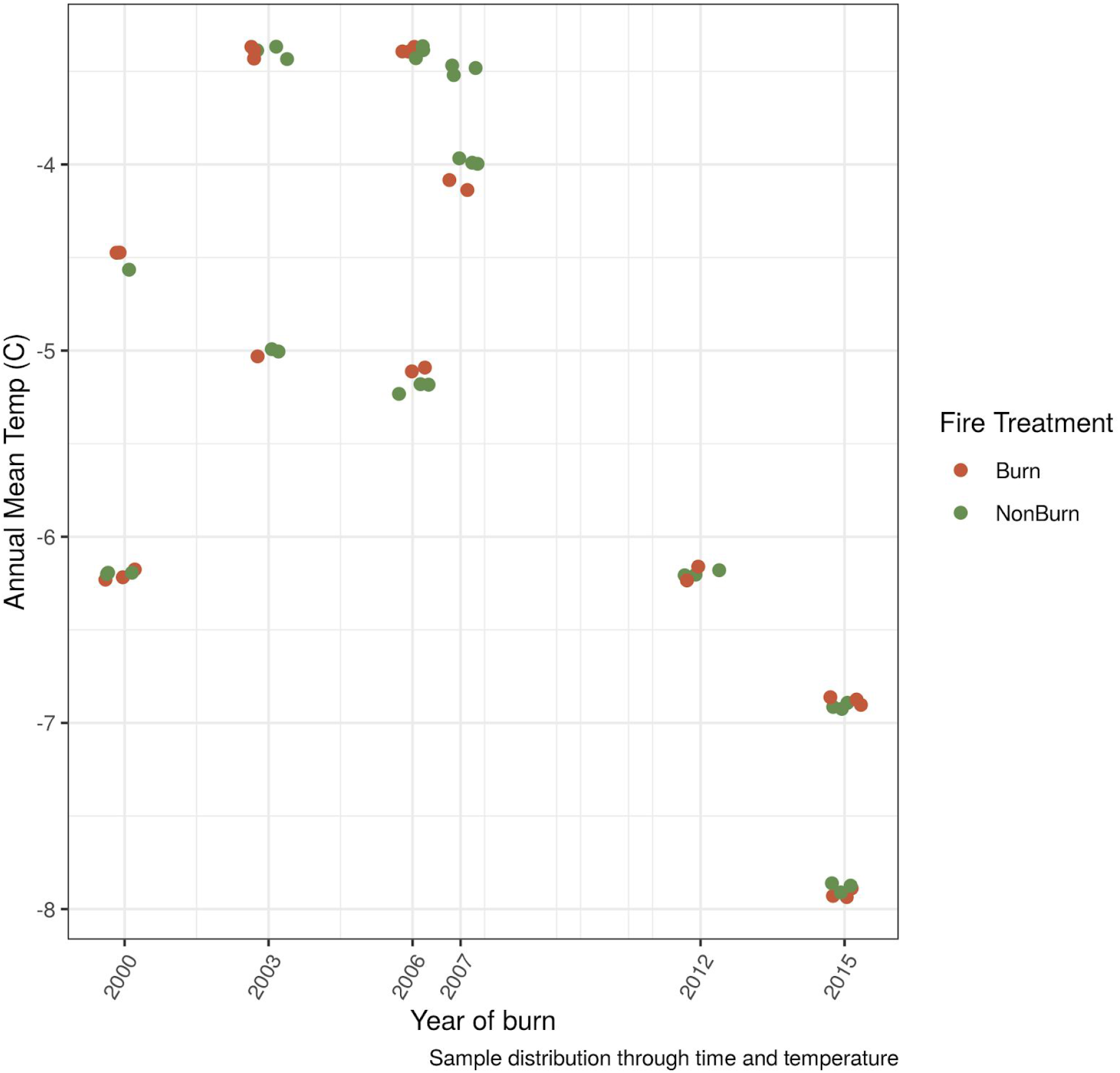
Sample distribution through time and temperature. Correlation between burn year and Mean Annual Temp could be due to climate change already impacting alpine areas over the past 20 years.

### Supporting Tables

#### Measured site characters

**For full metadata, see supporting data files** https://raw.githubusercontent.com/gzahn/Utah_Fires/master/Fire_metadata_plus_GDC.csv https://raw.githubusercontent.com/gzahn/Utah_Fires/master/plant_metadata_by_site.csv

**S.I. Table 1.**
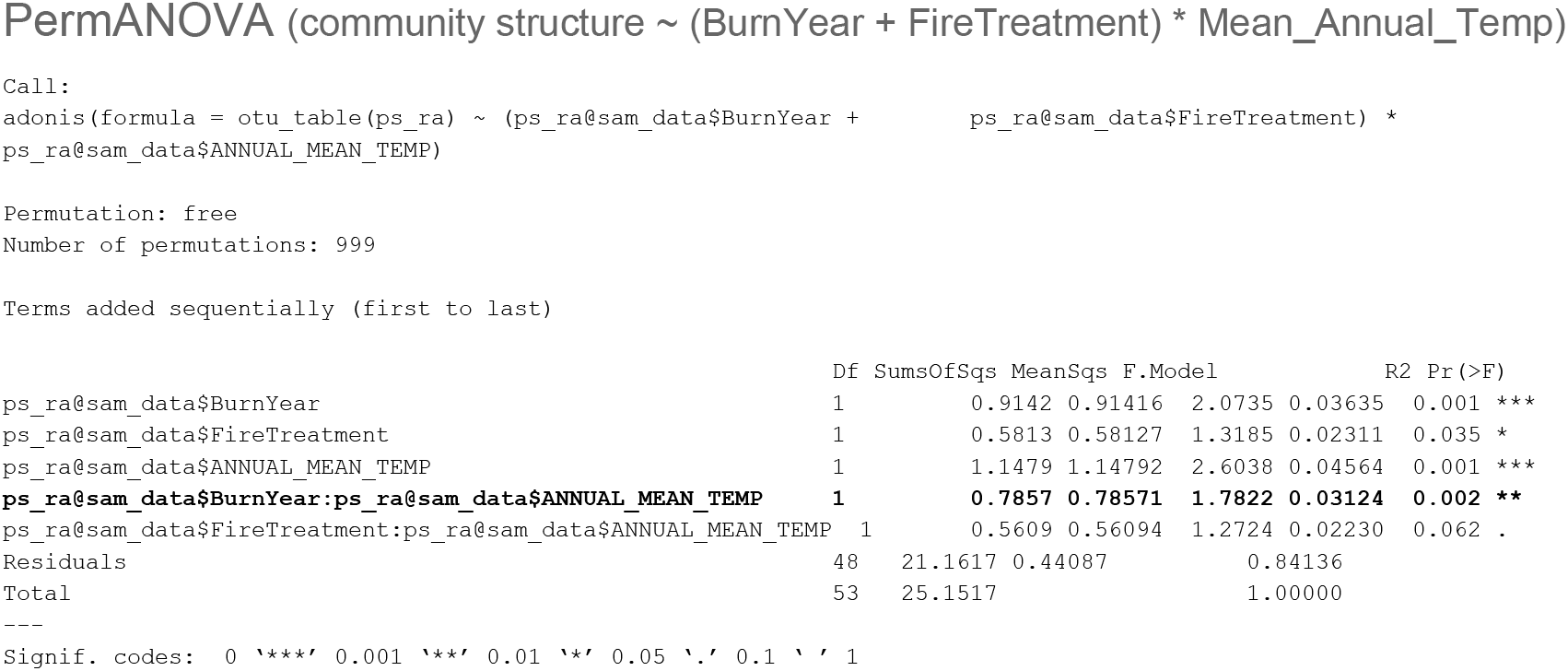
Permutational ANOVA shows significant interaction between Burn Year and Annual Mean Temperature for predicting community structure.

**S.I. Table 3.**
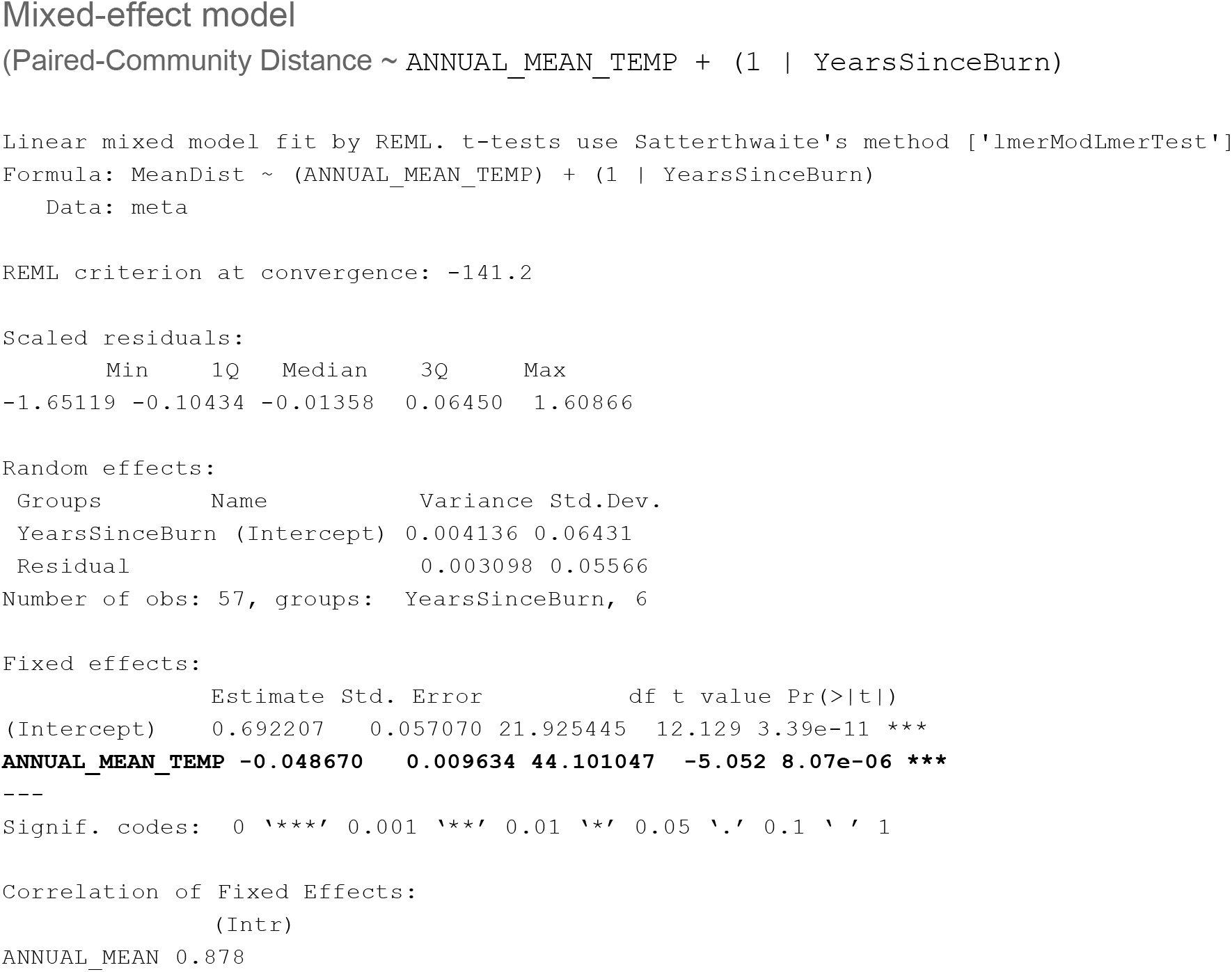
Mixed-effect model output

**S.I. Table 4.** Mantel test summary

| Std.Obs | Expectation | Variance |
| --- | --- | --- |
| 5.2980014858 | -0.0004410362 | 0.0017658801 |

